# Amlodipine as an Adjunct Bacterial Growth Inhibitor in Iron-Overloaded Conditions: In Vitro Effects Against *E. Coli*

**DOI:** 10.1101/2025.07.16.665167

**Authors:** Selim Mehmet Eke, Arnold Paul C. Cua

## Abstract

**Background:** Iron overload is a frequently underrecognized condition with a rising global prevalence. Excessive accumulation of iron has been considered a risk factor for the exacerbation of bacterial infections. Ca^2+^ is universally recognized as an indispensable cation for cell signaling. L-type voltage-dependent calcium channels (LVDCCs) facilitate the transport of Ca^2+^ and Fe^2+^ ions into the cytoplasm of eukaryotic cells. A non-protein molecule acting as a calcium channel has been found in prokaryotes. This finding has led to the proposal of a variety of signaling mechanisms for Ca^2+^ in bacteria. By combining research regarding the proliferation of bacteria exposed to iron and the potential role of Amlodipine not only as a mediator of iron overload at the cellular level but also as a growth inhibiting agent, we aim to test whether Amlodipine would have significant effects on the growth of *E. Coli* in iron-enriched microenvironments.

**Methods:** Ferrous sulfate was utilized to simulate an iron overloaded environment in-vitro by creating an agarose gel mixed with ferrous sulfate. *E. coli* was uniformly inoculated across the plates using standardized dilution protocol. A zone of inhibition test consisted of soaking sterile paper disks in increasing concentrations of Amlodipine (0, 1, 2, 3, 4, 5 mg/mL) and placing them on the agar. Zones of inhibition were measured, and results were statistically analyzed using one-way ANOVA for assessment of statistically significant differences between the means of the 6 groups and post-hoc Tukey HSD testing for comparison of individual groups to one another.

**Results:** Amlodipine exhibited a statistically significant, dose-dependent inhibition of *E. coli* growth (p = 6.1594 × 10 □ □). The diameter of the zones of inhibition increased consistently with Amlodipine concentration, with an R^2^ value of 0.9885, indicating a strong correlation. Higher concentrations of Amlodipine showed the greatest growth inhibiting effect, while the control group lacking Amlodipine exhibited little to no inhibition.

**Conclusion:** This study suggests that Amlodipine may inhibit bacterial growth in iron-overloaded conditions, highlighting its potential as an adjunct therapeutic option for patients with infections who are simultaneously at risk of iron overload. Further in vivo studies are warranted to assess its clinical relevance.

## 1 Introduction

Calcium (Ca^2+^) is a universally utilized signalling ion for eukaryotes (Clapham 2007). Ionized molecules, like the cation Ca^2+^, are hydrophilic and thus unable to pass through hydrophobic regions of the lipid bilayer (Kaas et al. 1989). Voltage-gated Ca^2+^ channels mediate the entry of extracellular Ca^2+^ into smooth muscle and cardiac myocytes and sinoatrial and atrioventricular nodal cells in response to electrical depolarization. Voltage-gated Ca^2+^ channels can be classified into the L, N, and T subtypes. Typically, the influx of Ca^2+^ via L-type voltage-dependent calcium channels initiates the process of vascular smooth muscle contraction (Bulsara et al. 2024). The Ca^2+^ then binds to intracellular calmodulin, which activates myosin light-chain kinase (MLCK). MLCK phosphorylates the myosin light chain, contracting smooth muscle and causing vasoconstriction, which is amplified by calcium-induced Ca^2+^ release from the sarcoplasmic reticulum. This chain of events results in a decreased vascular cross-sectional area, greater blood pressure and vascular resistance.

Resting cells in eukaryotes are known to maintain low levels of cytoplasmic Ca^2+^. Similarly, prokaryotes also maintain cytosolic Ca^2+^ levels well below that of the extracellular matrix (Gangola et al. 1987). Cellular organization can be altered by the influx of calcium ions in electricity-conducting cells, such as cardiomyocytes and neurons. Similar to sensory neurons in vertebrates, a voltage-induced calcium influx has been found to contribute to mechanosensation in *E. coli* (Bruni et al. 2017). Evidence has been found of a nonproteinaceous polymer functioning similarly to a voltage-gated calcium channel in *E. coli*, emphasizing a potential role for calcium signalling in *E. coli* (Das et al. 1997). Based on these findings, it is hypothesized that cytoplasmic calcium influx and voltage changes in *E. coli* promote ATP creation, cell division, and antibiotic resistance (Verstraeten et al. 2015). A correlation has been identified between the cytoplasmic calcium levels in bacterial cells and their pathogenicity and ability to spread (Hu et al. 2011). In *S. Pneumoniae,* it has been observed that increases in intracellular calcium concentration are able to result in oxidative defenses (Rosch et al. 2008). ATP-dependent transport is used by Streptococci for regulation of intracellular Ca^2+^ levels (Ambudkar et al. 1986). Wall-deficient forms of *E. Coli* necessitate calcium, likely via a cytoskeletal role (Onoda et al. 1999). Several potential signaling mechanisms of Ca^2+^ in *E. Coli* are demonstrated below in **Figure 1.**

**Figure 1.**
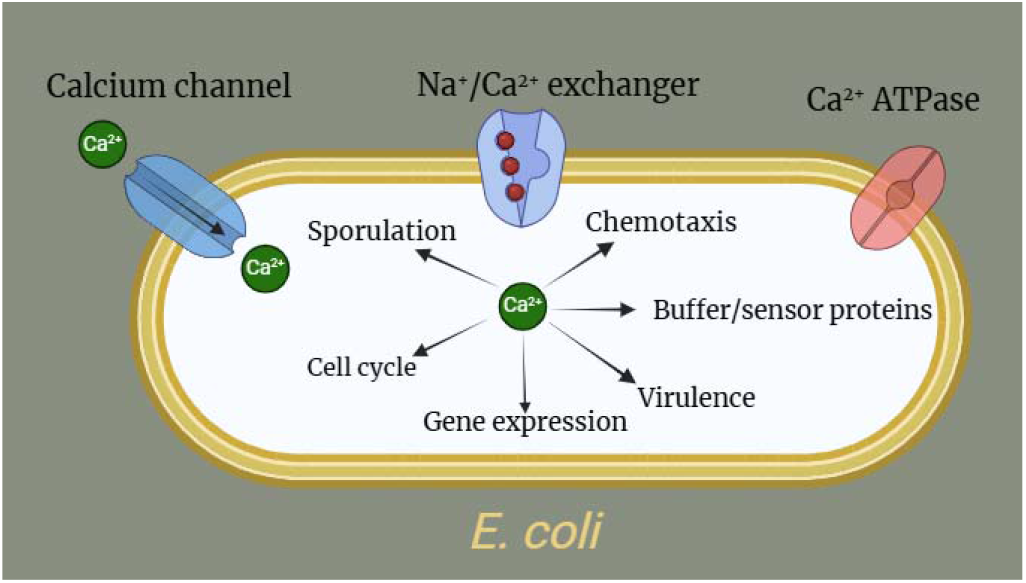
Proposed signaling mechanisms of Ca^2+^ resulting from its entry into *E. coli* bacterium via a nonproteinaceous ion channel and Na^+^/Ca^2+^ exchanger (Domínguez 2018).

Amlodipine (AML) is an oral dihydropyridine calcium channel blocker (Bulsara et al. 2024).

Approved by the Food and Drug Administration as Amlodipine Besylate in 1987, it has served as essential regulator of hypertension, chronic stable angina, vasospastic angina, and angiographically documented coronary artery disease (Bulsara et al. 2024). Calcium channel blockers produce their effects by binding to the 1 subunit of L-type voltage-gated Ca^2+^ channels and reducing Ca^2+^ flux through the channel. Dihydropyridines like Amlodipine and Nifedipine are known to exhibit vascular selectivity (Rysz et al. 2020). The primary side effects of Amlodipine include, but are not limited to, palpitations, edema, dizziness, and flushing. Dihydropyridine CCBs have an affinity for peripheral blood vessels and as a result are less likely to cause cardiac conduction abnormalities or myocardial depression compared to other CCBs. In addition, Amlodipine can serve as an antioxidant and upregulate the effect of nitric oxide, in turn working as a vasodilator (Mason et al. 2013). Reactive oxygen free radical species produced by non-transferrin-bound iron can be inhibited by Amlodipine (Crowe and Bartfay 2002).

Iron overload has climbed in prevalence in the past few decades due to the ease of access to iron and genetic disorders such as hereditary hemochromatosis, sickle cell anemia, thalassemia major, myelodysplastic syndrome, and sideroblastic anemia, yet the dangers of iron overload remain largely unrecognized. Though less discussed than iron deficiency, iron overload can silently wreak havoc on the body, leading to significant organ damage if left undetected. Excess iron, whether from frequent transfusions, intravenous iron administrations, or genetic conditions like hereditary hemochromatosis, accumulates in vital organs such as the liver, heart, and pancreas. Over time, this build-up can cause irreversible damage, including hepatic fibrosis, cardiac complications including cardiomyopathy, and dysfunction within the endocrine system such as bronze diabetes. As excess iron often goes unnoticed, early detection and targeted treatment strategies are crucial to prevent long-term damage.

Despite its established use as an antihypertensive, further research has demonstrated the surprising effects of Amlodipine in limiting myocardial damage, which occurs in the setting of iron overload. Past studies have explained the transport of myocardial iron via L-type voltage-dependent Ca^2+^ channels (LVDCCs). Treatment with calcium channel blockers (CCBs) has been shown to prevent deterioration of systolic and diastolic function, along with bradyarrhythmia, in iron-overloaded mice. Treatment with these CCBs also demonstrated a reduction in the composite index of myocardial damage and inflammation and the degree of apoptosis by reducing the potential for myocardial hemosiderosis. The effect of Amlodipine on mammalian calcium channels is depicted below in **Figure 2.**

**Figure 2.**
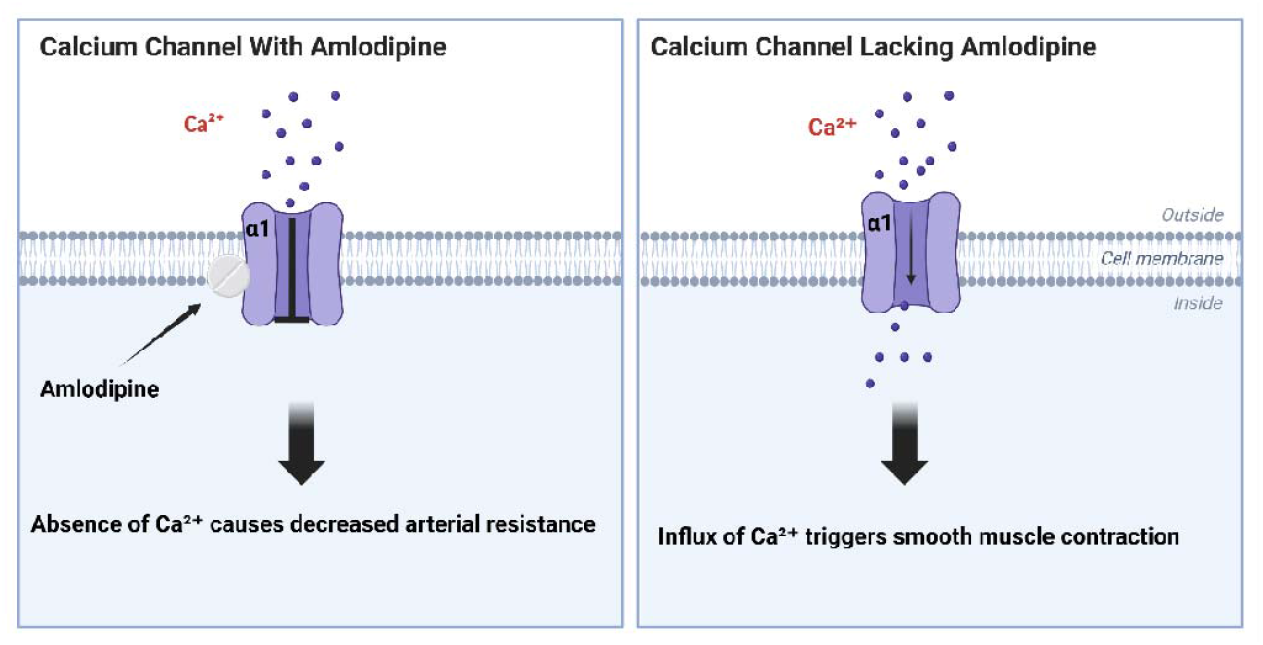
Comparison of the Effects of Ca^2+^ influx in a vascular cell membrane with versus without Amlodipine.

In addition to its well-recognized cardiovascular effects, Amlodipine has been identified by researchers to be a helper compound as a non-antibiotic which expresses growth inhibitory effects (Mazumdar et al. 2010). It is bactericidal towards *S. aureus* and *L. monocytogenes* (Mazumdar et al. 2010). Patterns of time-kill kinetics have confirmed Amlodipine’s bactericidal activities with a greater than 99.9% reduction in survival from the original inoculum (Akinjogunla et al. 2021). In gram-negative bacteria such as *Enterobacter spp., E. coli, K. pneumonia, and P. aeruginosa*, Amlodipine exhibited a concentration-dependent inhibition of bacterial growth (Akinjogunla et al. 2021). It has also been recognized as tissue-protective and anti-inflammatory against bacterial rhinosinusitis (Tatar et al. 2017).

Iron is considered an essential growth factor for bacteria, fungi, protozoans, and their vertebrate hosts (Weinberg 1978). For many years, it has been well-established that iron plays a significant role in bacterial metabolism (Hershko et al. 1988). Bacteria internalize Fe^2+^ by utilizing membrane-bound transferrin receptors and siderophores, which are high-affinity iron-binding proteins (Pieracci et al. 2005).

As amlodipine concentration increases, the zone of inhibition of *E. coli* bacteria is expected to increase. Because increased amlodipine concentration mediates iron accumulation, thereby inhibiting bacterial growth, this investigation is dedicated to determine whether Amlodipine has similar inhibitory effects on bacterial growth in iron-overloaded microenvironments (Oudit et al. 2003).

## Materials and Methods

*Escherichia coli* cultures were prepared following the standardized protocol developed by Son and Taylor with minor modifications (Son and Taylor 2012). Following incubation, visible growth from the streaked area was used to inoculate 5 mL of sterile Luria Broth without selection for single colonies. The broth cultures were incubated at 37°C with shaking (aerobic conditions) for 16–18 hours to reach a high-density bacterial suspension, which was used for subsequent experimental assays.

5 grams of Luria Broth Agar were weighed and placed into a 250 mL Erlen-Meyer Flask. 200 mL of distilled water was added to the solution and mixed. 3.45 mL (3,450 µL) of ferrous sulfate were micropipetted into the solution and mixed once more. This quantity was determined based on a standardized calculation to simulate an iron overloaded microenvironment in-vitro, necessitating 1000 µM per liter of iron within the Agarose gel (Landrigan et al. 1989). A 200 mL solution would thus require 200 µM of iron. Hence, the calculation is as following:

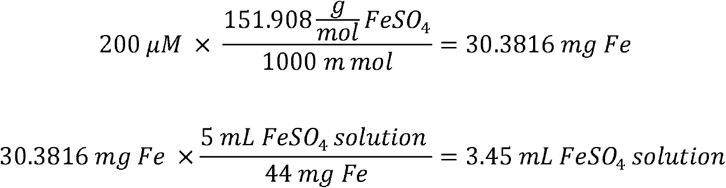

The mixture was microwaved for 1 minute and 30 seconds. The Agarose gel containing ferrous sulfate was allowed to cool for 15–20 minutes before being poured into each of the six plates. The gel was left to harden in the plates for 10 minutes. Bacteria were inoculated using an inoculation loop sterilized via a Bunsen burner. Stock *E. coli* was diluted in a 10 mL tube before 10 µL of the *E. coli* solution were micropipetted onto each of the six plates. A flat spreader was sterilized with isopropyl alcohol before being used to spread *E. coli* homogeneously across the plates. Six 200 mL beakers and the respective quantities of Amlodipine (0, 10, 20, 30, 40, and 50 mg) were gathered. 10 mL of water were added to each beaker and the solutions were mixed. 30 sterile paper discs were gathered, and five were immersed in each of the six solutions. Tweezers were sterilized with isopropyl alcohol and used to remove discs from the solutions and place them in the four corners and center of the Petri dishes. All Petri dishes were incubated for 24 hours at 37 □. After 24 hours of incubation, zones of inhibition were measured.

**Table 1.** Raw data for Amlodipine concentration (mg/mL) versus inhibition zone diameter of *E. coli* (∓0.05cm) for each group.

| Amlodipine concentration<br>mg/mL | Disk Number |  |  |  |  |
| --- | --- | --- | --- | --- | --- |
|  | 1 | 2 | 3 | 4 | 5 |
| | Inhibition Zone Diameter of *E. coli* ($\pm 0.05$cm) | | | | |
| Control | 0.00 | 0.00 | 0.00 | 0.00 | 0.00 |
| 1 | 0.00 | 1.20 | 0.00 | 1.10 | 0.90 |
| 2 | 1.60 | 1.00 | 1.40 | 1.20 | 0.0 |
| 3 | 1.40 | 1.50 | 1.4 | 1.80 | 1.70 |
| 4 | 1.80 | 1.80 | 2.00 | 1.90 | 1.60 |
| 5 | 2.10 | 2.30 | 2.40 | 2.40 | 2.30 |

**Table 2.** Processed data for Amlodipine concentration (mg/mL) versus inhibition zone diameter of *E. coli* (∓0.05cm), in addition to standard deviation [σ =√(∑x−x□)^2^ / n] and the coefficient of variation [CV = (σ / x□) x 100] for each group.

| Amlodipine concentration<br>mg/mL | Mean Value Inhibition Zone Diameter of *E. coli* Bacteria in cm ($\pm 0.05$cm) | Standard Deviation | Coefficient of Variation |
| --- | --- | --- | --- |
| Control | 0 | 0 | 0 |
| 1 | 0.64 | 0.53 | 83.03 |
| 2 | 1.04 | 0.56 | 53.57 |
| 3 | 1.56 | 0.16 | 10.42 |
| 4 | 1.82 | 0.13 | 7.29 |
| 5 | 2.30 | 0.11 | 4.76 |

**Table 3.**
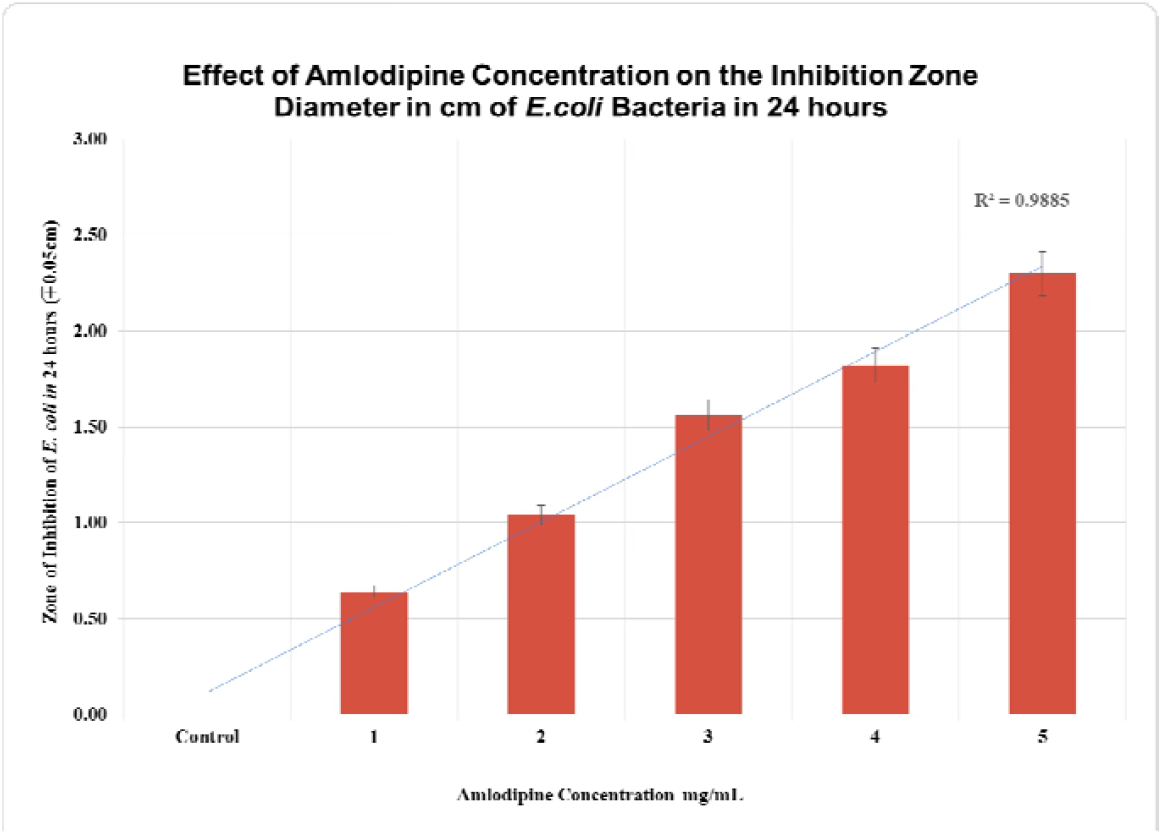
The effect of Amlodipine concentration on the inhibition zone diameters of *E. coli* bacteria after 24 hours of incubation.

## Results

This investigation aimed to understand whether Amlodipine had a growth inhibiting effect on *E. coli* in iron-enriched microenvironments. To determine if there is a statistically significant difference between the means of any of the six treatment groups, a One-Way ANOVA (Analysis of Variance) was the optimal choice (Ross 2017). In Table 9, the treatment Sum of Squares (SS) represents the variability between the six treatment groups, 17.4947. The Error Sum of Squares indicates the variability within each treatment group, 3.2440. The total Sum of Squares represents the total variation in the dataset, a sum of Treatment Sum of Squares and Error Sum of Squares, 20.7387.

Treatment Degrees of Freedom (*df*) was calculated as 6 treatments – 1, hence *df = 5*. Error *df* was the total number of data points, 30, minus the number of groups, 6. Hence, the error *df* = 24. Total *df* is the total number of observations (30) minus 1, so total *df* = 29. The mean square (MS) represents the average variation within and between groups. Treatment MS is treatment SS (17.4947) / treatment *df* (5), hence treatment MS = 3.4989. Error MS is error SS (3.2440) / error *df* (24), hence error MS = 0.1352. The F-Statistic of 25.8861 was calculated as Treatment MS / Error MS, or 3.4989/0.1352=25.8861, meaning that the variability between group means was 25.8861 times larger than the variability within the groups. A very large F-Statistic indicates that the differences between group means are significantly larger than the variability within each group. As a result, it can be claimed that at least one group’s mean is significantly different from the others. Additionally, a very small p-value of 6.1594 × 10^−9^ is below 0.05, aligning with the interpretation of the F-Statistic of very strong statistically significant differences between the means of at least one of the six groups. Therefore, it can be concluded that treatment with varying concentrations of Amlodipine on *E. col* in an iron-enriched microenvironment had statistically significant effects between at least one of the six groups.

As shown in Graph 1, an R^2^ value of 0.9885 indicates a high correlation between the independent variable and the dependent variable. Thus, 98.85% of the variability in the dependent variable can be explained by the variability in the independent variable. This confirms the relationship between Amlodipine concentration and the inhibition zone diameter, and the degree to which the variation in bacterial inhibition can be attributed to increasing Amlodipine concentration. A high R^2^ value strengthens the hypothesis by demonstrating the replicability of Amlodipine’s growth inhibitory activity against *E. coli*. As shown by the data, the non-overlapping of error bars shows the stability of the zone of inhibition bacterial values at different Amlodipine concentrations. Standard deviation values were also calculated for each mean inhibition zone diameter. The results of the study also showed that higher Amlodipine concentrations led to greater inhibition of *E. coli* bacterial growth.

The post-hoc Tukey HSD Test depicted in **Table 4** was performed for the six treatments (Kim 2015). Multiple pairs of conditions (A-F) were tested to see if there were statistically significant differences between their means. A/E, A/F, B/D, B/E, B/F, and C/F pairs were found to have strongly significant differences in their means, with p < 0.01 in each of the six pairs. C/E and D/F had moderately significant differences between their means, with p < 0.05 in each of the two pairs. B/C, C/D, D/E, and E/F did not have significant differences between their means, with p > 0.05 for each of the four pairs. As groups E and F exhibited significant inhibition when compared to groups A and B, it can be claimed that Amlodipine had a concentration-dependent growth inhibitory effect on E. coli. The highest Amlodipine concentrations, at 4 mg/mL and 5 mg/mL for E and F, respectively, therefore resulted in the greatest inhibition of growth according to the post-hoc Tukey HSD Test.

**Table 4.** Post-hoc Tukey HSD Test for comparison between individual group means of Amlodipine concentration versus inhibition zone of sterile paper disks.

| treatments pair | Tukey HSD Q statistic | Tukey HSD p-value | Tukey HSD inference |
| --- | --- | --- | --- |
| A vs B | 3.8925 | 0.1010949 | insignificant |
| A vs C | 6.3253 | 0.0019641 | ** p<0.01 |
| A vs D | 9.4880 | 0.0010053 | ** p<0.01 |
| A vs E | 11.0693 | 0.0010053 | ** p<0.01 |
| A vs F | 13.9887 | 0.0010053 | ** p<0.01 |
| B vs C | 2.4328 | 0.5285981 | insignificant |
| B vs D | 5.5955 | 0.0068815 | ** p<0.01 |
| B vs E | 7.1768 | 0.0010053 | ** p<0.01 |
| B vs F | 10.0962 | 0.0010053 | ** p<0.01 |
| C vs D | 3.1627 | 0.2585426 | insignificant |
| C vs E | 4.7440 | 0.0280382 | * p<0.05 |
| C vs F | 7.6634 | 0.0010053 | ** p<0.01 |
| D vs E | 1.5813 | 0.8558976 | insignificant |
| D vs F | 4.5007 | 0.0411044 | * p<0.05 |
| E vs F | 2.9194 | 0.3382499 | insignificant |

The coefficient of variation (CV) decreases with the increase in concentration of Amlodipine, which indicates that the accuracy of inhibition zone measurements is larger at higher concentrations of Amlodipine. The lowest CV of 83.03% was observed at 1 mg/mL, which indicates greater variability in bacterial response at lower concentrations.

These results support the research hypothesis which states that there is a dose-dependent growth inhibitory effect of Amlodipine. The results may have important clinical relevance and could support the development of innovative approaches to antimicrobial therapy.

## Discussion

This study found that Amlodipine may present a concentration-dependent inhibitory effect on *E. coli* growth within an in-vitro, iron-overloaded microenvironment. Fe^2+^ is known to compete with Ca^2+^ for the C-terminal cytoplasmic Ca^2+^ binding site (Bers 2002). LVDCCs are believed to be the only Fe^2+^ transporter with their uniquely increasing activity in settings of elevated iron concentrations (Randell et al. 1994). There is evidence of a link between LVDCCs and myocardial iron deposition. In mice studies, Amlodipine was proven to reduce cardiac complications protecting diastolic and systolic dysfunction (Oudit et al. 2003). *E. coli* presents mechanosensitive channels, which function through a nonproteinaceous polymer acting as a voltage-gated calcium channel (Bruni et al. 2017). Voltage depolarization allows for calcium influx into the intracellular space of *E. coli.* This influx is believed to contribute to the cellular functioning of prokaryotic organisms. In summary, this data along with past findings indicate the potential for Ca^2+^ as a signaling mechanism for pathogenicity in *E. coli*.

## Conclusion

Our findings indicate that Amlodipine presents a mechanism for inhibition of bacterial growth in iron-enriched settings. Amlodipine exhibited a dose-dependent zone of inhibition when using sterile paper disks and varying concentrations of Amlodipine. Significant differences between the control group and the highest Amlodipine concentrations allow us to presume a steady increase in inhibition zones as Amlodipine dosage is increased.

The pharmacological inhibition of voltage-gated calcium channels has been studied in rodents so far and has been found to present cardioprotective effects, and there may be an even greater therapeutic effect in human beings in the setting of iron overload and aggressive bacterial infections. As humans do not have an excretory mechanism for iron, blockage of calcium channels may be a novel way to prevent excess iron accumulation. Since this study remains in-vitro, further research is necessary to determine whether Amlodipine has similar effects in iron overload settings in humans with bacterial infections.

## Data Availability

The data are contained within the article.

## Author Contributions

Writing – original draft preparation, methodology, statistical analysis, S.M.E; writing – review and editing, S.M.E. and A.C. All authors have read and agreed to the final version of the manuscript.

## Acknowledgments

We would like to thank our team for their dedication and hard work during study design, experimentation, and creation of this manuscript.

